# Reproductive plasticity in both sexes interacts to determine mating behaviour and fecundity

**DOI:** 10.1101/2021.02.11.430788

**Authors:** E.K. Fowler, S. Leigh, A. Bretman, T. Chapman

## Abstract

Organisms alter their phenotype in response to variation in their environment by expressing phenotypic plasticity. Both sexes exhibit such plasticity in response to contrasting environmental and social cues, and this can reflect the influence of sexual conflict. However, theory predicts that plasticity expressed by both sexes may *either* maximise the sex-specific fitness of both, or of one sex at the expense of the other. Hence empirical tests of the predictions are sorely needed. Here we conducted novel tests of the fitness effects of interacting reproductive plasticity in *Drosophila melanogaster*. First, prior to mating, males were kept alone, or with same sex rivals, and females were kept alone, in same sex, or mixed sex groups. Second, we conducted matings between individuals from all these social treatments under ‘choice’ and ‘no choice’ scenarios. The results showed that males and females can both plastically respond to these socio-sexual environments to influence the expression of mating duration, mating latency, and fecundity. These plastic responses interacted significantly to determine mating latency and fecundity. Effects on mating latency were also observed under both choice and no-choice conditions, but in opposing directions. Variation in the outcome of interacting plasticity pivoted around the outcomes observed with focal females that had been maintained in same-sex environments prior to mating. However, not all fitness-related traits examined responded in the same way. Mating duration was determined largely by the social environment of the male. Our results show that the expression of some, but not all fitness-related reproductive traits can be determined by the outcome of interacting behavioural plasticity expressed by both sexes. This highlights the need for new predictive theory informed by these empirically-derived parameters. Overall, we conclude that variation in the expression of shared traits due to interacting plasticity represents an important and novel facet of sexual interactions.

**Impact Summary:** Animals and plants are able to respond to variation in their environment by expressing phenotypic plasticity. In sexual organisms, both males and females can exhibit such plasticity but the cues they respond to and the fitness consequences of these actions may be different between the sexes, and even conflicting. For example, males may respond to the presence of competitors by altering their mating behaviour or ejaculate transfer to increase their own, but not necessarily their mate’s reproductive output. However, females may also express phenotypic plasticity in response to their social and sexual environment to maximise their own fitness. Theory suggests that plasticity expressed by both sexes may either maximise the sex-specific fitness of both, or of one sex at the expense of the other. So far, little experimental work has been conducted to explore such interacting plasticity. Here we conducted novel tests of the fitness effects of interacting plasticity in the fruit fly *Drosophila melanogaster*. In doing so, we provide novel experimental evidence for interacting behavioural plasticity. We show that males and females can plastically respond to their socio-sexual environment to influence the expression of mating duration, mating latency, and fecundity. These plastic responses, while induced to increase the fitness interests of each sex, interact in the case of mating latency and fecundity and may reflect the outcome of sexual conflict. Our findings suggest that studies of reproductive behaviour should carefully consider the socio-sexual environment of both males and females and highlight the need for new predictive theory informed by empirically-derived parameters. Overall, we show that interacting plasticity between sexes represents an important and novel facet of sexual interactions.

## Introduction

Phenotypic plasticity is the ability of organisms to directly respond to biotic or abiotic changes in the environment by altering their phenotype (Komers et al., 1997; West-Eberhard, 2003; Fordyce, 2006; Dingemanse and Wolf, 2013). Potential benefits of plasticity lie in maximising fitness or survival in variable environments in which a fixed strategy may be costly (Bretman et al., 2013a). Therefore, strongly plastic individuals may be better able to match the conditions experienced in a heterogenous environment. The socio-sexual environment is an important stimulus for the expression of behavioural plasticity across different taxa (e.g. Han and Brooks, 2014; Dorset et al., 2017; Oku and van den Beuken, 2017). For example, information such as the number of rivals or mating opportunities can indicate to an individual the likely level of competition for resources and mates they will face (Davis et al., 2011; Bretman et al., 2011).

Behavioural plasticity has been well studied in fruitflies. Male *D. melanogaster* are observed to increase aggressive behaviour towards rival males when the number of rivals is low but decrease aggression when high (Nandy et al., 2016). This results in an increased chance of mating when competition is less likely, and avoids potentially damaging conflict or diminishing fitness returns when competition is strong (Nandy et al., 2016). *D. melanogaster* males may also alter their reproductive investment in response to perceived levels of sperm competition. For example, males exposed to rivals prior to mating extend mating duration and transfer into females more of two key seminal fluid proteins (SFPs), Ovulin and sex peptide (SP) (Wigby et al., 2009; Bretman et al., 2013a; Filice et al., 2020). Ovulin and SP induce important post-mating behavioural and physiological changes to females, including increased fecundity and decreased sexual receptivity (Chapman et al., 1993; Herndon and Wolfner, 1995; Heifetz et al., 2000; Liu and Kubli, 2003; Wigby et al., 2009). This is expected to benefit males as they invest in energetically costly SFPs that increase fecundity and decrease female re-mating receptivity only when required (Wigby et al., 2009). As the receipt of SFPs can be costly for females (Chapman et al., 1995; Wigby and Chapman, 2005) this may also exacerbate sexual conflict (Sirot et al., 2015).

Though understudied outside the context of mate choice, there are increasing reports of female plasticity. For example, females can observe and learn oviposition strategies from other females, choosing to lay eggs on a potentially ‘good’ food substrate (Sarin and Dukas, 2009). Mated females may also alter their egg laying behaviour in response to a male-derived pheromone, aggregating in order to lay eggs (Wertheim et al., 2006), and can also exhibit variation in fecundity according to the genetic diversity of the males in their social environment (Billeter et al., 2012). The intrasexual pre-mating environment can influence a female’s behavioural plasticity. For example, female *D. melanogaster* housed with other females lay significantly more unfertilised eggs as virgins but are less fecund following mating, compared to socially isolated females (Fowler et al., 2020; Churchill et al., 2021).

Though increasing evidence shows that both sexes can express reproductive plasticity in response to the presence of conspecifics, we lack information on whether plastic responses can interact in determining the overall levels of reproductive investment made by each sex. For example, we do not yet know how the expression of plastic responses by one sex affects those of the other. This important omission is the main focus of the current study. Interactions between plasticity expressed by males and females are expected to be an important determinant of overall fitness. For example, we hypothesise that the plastic response of a male could trigger females to alter their own reproductive investment. However, we lack theory on which to base predictions, and that which does exist predicts variable outcomes. For example, interacting plasticity is predicted by theory to either maximise the sex-specific fitness of *both* sexes (McGhee et al., 2013), *or* of one at the expense of the other (Yamaguchi and Iwaga, 2015; McLeod and Day, 2017; Day and McLeod, 2018). Variation in outcomes might be predicted if the strength of sex-specific selection on the relevant fitness traits involved differs, or if one sex has gained the upper hand in the expression of traits subject to sexual conflict. However, to our knowledge, there are as yet no empirical tests of this theory or of the effects of interacting plasticity on mating behaviour and fitness. Here we addressed this omission by using the fruitfly *Drosophila melanogaster* to conduct novel tests of the effects on key fitness traits of plasticity expressed by both sexes in response to their social and sexual environments.

We first varied the pre-mating social environments, by keeping focal flies alone, with same-sex or opposite sex individuals. We predicted that focal females exposed to other females, vs to males would vary in their perception of resource vs sexual competition (Table 1). Similarly, that males exposed to rivals would perceive a high threat of sexual competition. We then mated males and females from each of the social environments varying in competitiveness together in all combinations. This allowed us to test for interacting effects of plastic response of both sexes to those environments on mating success, mating latency, mating duration and fecundity.

**Table 1.** Description of the pre- and post-mating treatments and mating environments used in this study. The environments and the expected effects of those environments are described.

| Treatment |  | Environment description | Expected effect of treatment |
| --- | --- | --- | --- |
| **Females, pre-mating** | Alone | One focal female per vial. | Signals absence of competition. |
|  | Group same sex (GSS) | One focal female and three non-focal females per vial. | Signals competition between females for food, oviposition sites, and potentially males, is likely. |
|  | Group mixed sex (GMS) | One focal female and three males per vial. | Signals females likely to experience effects of competition between males for matings and fertilisations. |
| **Males, pre-mating** | No rival | One focal male per vial. | Signals low competition for matings and fertilisations. |
|  | Rival | One focal male and three non-focal males per vial. | Signals competition for matings and fertilisations likely. |
| **Mating arena** | Choice | One focal female and two focal males in the mating arena. One male is from the no-rival treatment and one from the rival treatment. | Females can simultaneously assess different males, who can directly compete. Assessment of competition based on previous and current experience of both sexes. |
|  | No choice | One focal female and one focal male in the mating arena. | No opportunity for direct comparisons. Assessment of competition, and choice of whether to mate at all, is indirect and based upon previous experience only. |

We conducted mating tests in both traditional ‘choice’ and ‘no-choice’ mating scenarios. The choice assays placed females from each of three social treatments (alone, same-sex or opposite-sex) with two males from two different social treatments (alone with no rival, or with same-sex rivals). This introduced the effect of direct competition between males of different environments and of direct comparison of those males by females (Table 1). In the no-choice assays, females from each of the three social treatments were allocated a single male from either social treatment. This removed the effect of direct competition and tested the effect of responses to previous social environments (Table 1).

## Methods

### Stock Maintenance and Fly Collection

Wild type *D. melanogaster* flies were from a large laboratory population originally collected in the 1970s in Dahomey (Benin). Flies were maintained in stock cages with overlapping generations on SYA medium (Sugar Yeast Agar: 100 g brewer’s yeast, 50 g sugar, 15 g agar, 30 ml Nipagin (10% w/v solution), and 3 ml propionic acid, per litre of medium). SYA was used throughout the experiments and all flies were cultured and reared, and all experiments performed, at 25°C, 50%RH, on a 12h:12h light:dark cycle. Eggs for all experimental manipulations were collected from population cages using purple agar egg collection plates (275ml H_2_O, 12.5g agar, 250ml red grape juice, 10.5ml 10% w/v Nipagin solution) supplemented with live yeast paste. First instar larvae were picked into 7ml SYA vials (75 x 25 mm) at a density of 100 larvae per vial. Adults were collected within 8h of eclosion, separated into same sex groups using ice anaesthesia and stored 10 per vial. Adults were stored under these conditions for four days and allowed to reach sexual maturity until use in the experiments.

### Manipulation of Focal Female and Male Pre-Mating Social and Sexual Environments

*Females*: The three female social environment treatments were: ‘alone’, ‘group same sex’ (GSS), and ‘group mixed sex’ (GMS). The female social environment treatments were all set up in vials that were divided into two chambers using perforated acetate (through which sound, smell and visual cues could be transmitted, but flies could not physically pass). For the alone treatment, a single focal female was placed in one of the vial chambers. For the GSS treatment, one focal female was placed in one chamber and three non-focal females in the other. For the GMS treatment one focal female was placed in one chamber and three non-focal males in the other. The plastic dividers limit physical contact between individuals, but we have found previously that pre-conditioning vials by allowing non-focal flies access to the entire vial for 24h prior to adding the focal female and dividers provides sufficient cues for focal females to accurately detect their social environment (Fowler et al., 2020). We therefore preconditioned the vials in the GSS and GMS treatments (figure 1A – ‘Vial Pre-conditioning’) by placing all non-focal individuals into the appropriate vial 24h before the introduction of the focal female and the acetate dividers.

**Figure 1.**
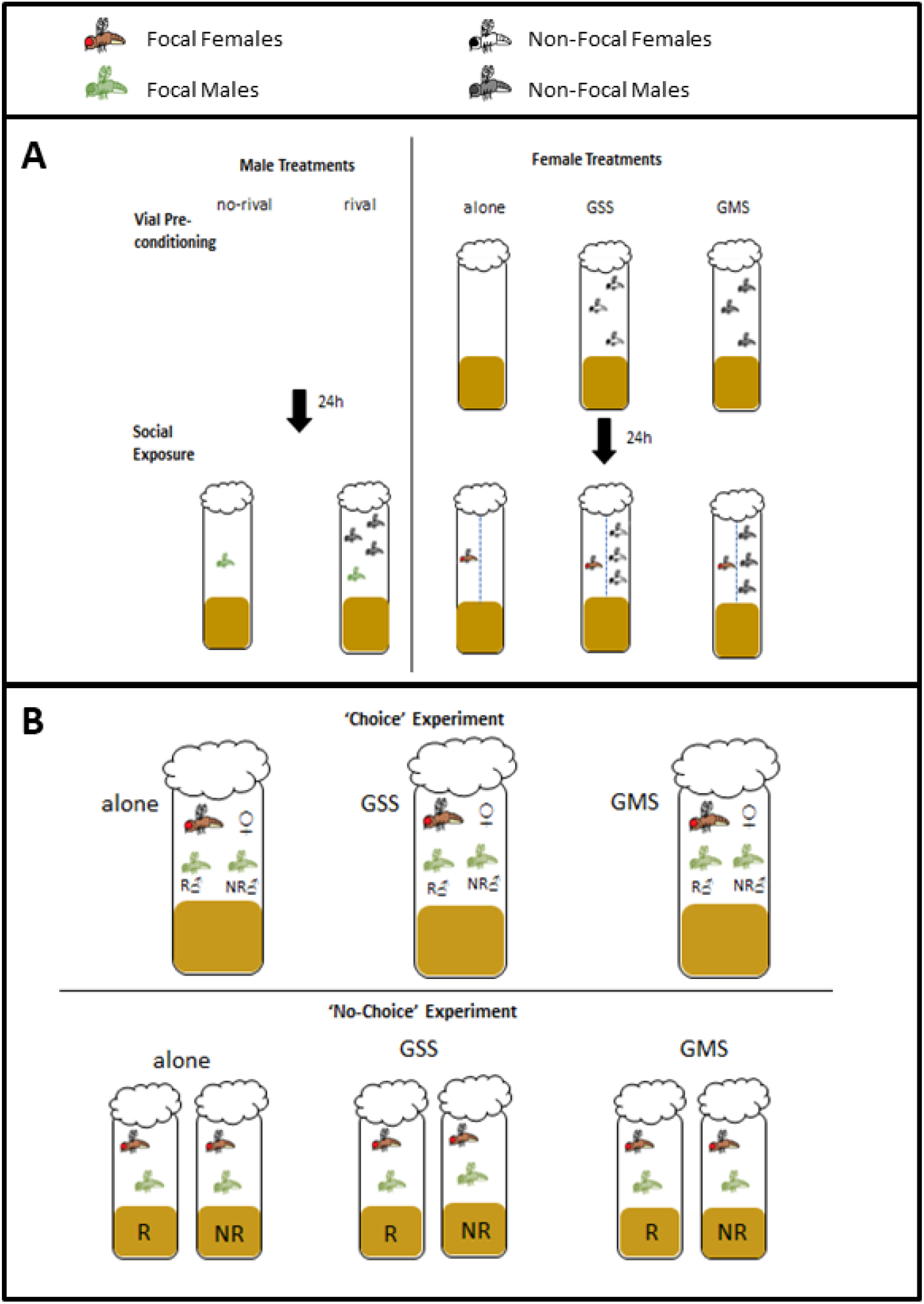
Pre- and post-mating environment manipulations. **A: Set up for the pre-mating social environment manipulations.** In the female treatments, non-focal flies were placed into vials 24h before the introduction of the focal female in order to transfer residual social cues in the group same sex (GSS) and group mixed sex (GMS) treatments (vial pre-conditioning). After 24h, pre-conditioned GSS and GMS and alone treatment vials were divided using perforated acetate sheets (blue dashed line) to separate focal females from the non-focal flies. **1B: Mating assay set up for the choice and no-choice experiments**. Vials in the choice experiment consisted of a single female from one of the three social treatments in a vial with both a rival (R) and no-rival (NR) male. The no-choice experiment consisted of a single female from one of the three social treatments in a vial with either a rival or no-rival male.

*Males*: The two male social environment treatments were: ‘rival’ and ‘no-riva’. The rival treatment consisted of four males per vial, and the no-rival treatment a single male in a vial. Male social treatments did not require pre-conditioning and so focal males were randomly allocated into one of the two male treatments at the same time as the focal females.

All females and males were kept in the above pre-mating social environments for 48h before the mating assays (figure 1A - ‘Social Exposure’). We conducted matings between the males and females kept in all these social environments in two separate experiments that varied only in whether they allowed for direct competition / choice. In the first, the mating test was conducted using a traditional ‘choice’ design, whereby each focal female was exposed to two males in the mating arena (one male from each of the rival or no rival social treatments). In the second experiment, mating tests consisted of one female and one male in a traditional ‘no-choice’ design (Table 1).

#### 1: Effect of male and female social environment on mating behaviour and fecundity under choice conditions

In the choice experiment, the mating and fecundity responses of alone, GSS and GMS females paired in the mating arena with two males, one from the rival and one from the no-rival treatments, were recorded. Males were wing marked with either red or black ink using Staedtler Lumocolor red/black marker pens. Wing marking was balanced across rival and no-rival groups to ensure that any effect on female choice of ink colour would not introduce any directional bias. On the day of the mating experiment, the focal female from each treatment was aspirated into a vial with marked males from the rival and no-rival treatments and given 90 minutes to mate (figure 1B – ‘Choice Experiment’). Mating latency and mating duration were recorded as was the ink colour of the male that was chosen / secured the mating. After mating had finished, males were discarded and females were left in the vials for 24h to lay eggs. The sample sizes for each mate choice scenario were ~50 per female treatment. Since only one male out of the pair secured a mating, samples sizes for mating latency, duration and fecundity were approximately 25 per male and female treatment, depending on how many matings occurred (see table S1 for final sample sizes).

#### 2: Effect of male and female social environment on mating behaviour and fecundity under no-choice conditions

In the no-choice experiment, the mating and fecundity responses of alone, GSS and GMS females placed with either rival or no-rival condition males were recorded. Focal females and males were in set up in their alone, GSS and GMS social environments, as described above, for 48h. In this set up females could mate or not mate with a rival or no-rival male assigned at random to them, thus marking of focal males was not required. All non-focal males were wing clipped under CO_2_ anaesthesia 24h prior to setting up the social exposure treatments. On the day of the mating experiment, focal females from each alone, GSS and GMS treatment were each aspirated into an SYA vial with one focal male either from the rival or no-rival treatment. Pairs were given 90 minutes to mate during which mating latency and duration were recorded (figure 1B – ‘No-Choice Experiment’). Post-mating egg data were collected as before. The starting sample sizes for mating latency, duration and fecundity were approximately 50 per female and male treatment (final sample sizes for each treatment are given in table S1.

### Statistical Analyses

All analyses were carried out using R version 3.6.3 (R Core Team, 2020) (packages used: survminer, survival, ggplot2, dplyr, stats). A Generalized Linear Model (GLM) with a quasi-Poisson error block was used to analyse post-mating egg production and mating duration for the rival and no-rival groups for each alone, GSS and GMS female social treatments. In these models, each female group was subset and tested separately for their response to the rival/no rival treatment. A Cox Proportional Hazards Model was used to test for differences in mating latency. Individuals that did not mate within 90 min were treated as censors. A Chi Square analysis was used to test for the effect of wing colour on male mating success. A GLM with binomial errors was used to test effect of male and female social treatment on proportion of matings secured. A maximal model was fitted, and then simplified to remove the male:female treatment interaction term. Models were compared using analysis of deviance (anova() in “stats” package) with a chi-squared test. The final model retained the male and female social treatment terms (see table S1 for final sample sizes).

## Results

### 1: Effect of male and female social environment on mating behaviour and fecundity under choice conditions

Each female was presented with males from rival or no-rival social treatments simultaneously, allowing females to directly compare different males and allowing males to compete. Thus intersexual choice and intrasexual competition was possible. Wing marking colour had no effect on male mating success within any of the female treatments (table 2). There was a marginally non-significant trend for no-rival males to secure more matings than rival males in each of the alone, GSS and GMS female social treatments (X^2^ = 3.4, *df =* (1,4), *p =* 0.06; table 2, figure 2). Matings occurred rapidly, with a median latency to mating of three minutes across all treatments. The social environment of the males had no significant effect in the GMS (*HR* = 0.861, *95% CI* [0.462, 1.604], *p =* 0.637) and alone (*HR =* 0.841, *95% CI* [0.466, 1.518], *p =* 0.566) treatments. However, in the GSS treatment, rival males started mating significantly faster than no-rival males (*HR =* 2.384, *95% CI* [1.232, 4.612], *p =* 0.0099) (figure 3). Females in the alone and GMS treatments mated for significantly longer with rival than with no-rival males (alone: *t_44_ =* 2.486, *p =* 0.0168; GMS: *t_41_ =* 2.799, *p = 0.0078*). The trend was in the same direction, but was not significant in the GSS treatment (*t_39_ =* 0.791, *p =* 0.434) (figure 4A). There was no significant difference in the number of eggs produced 24h after mating between females that mated with rival male and no-rival males in any of the female social environments (alone: *t_44_ =* 1.517, *p =* 0.136; GSS: *t_39_ =* −0.77, *p =* 0.446; GMS: *t_41_ =* −0.429, *p =* 0.67) (figure 5A).

**Table 2.**
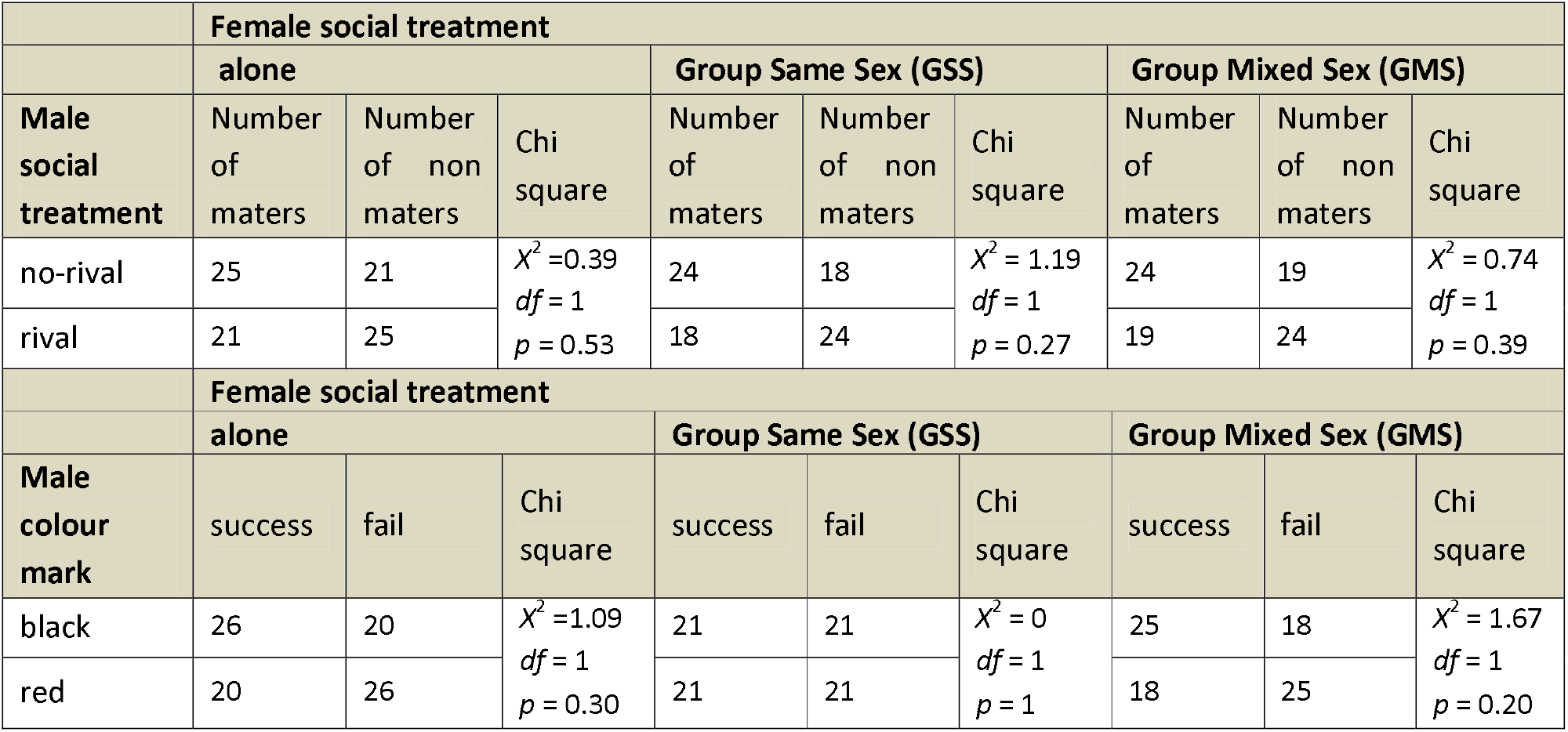
Number of matings by male social treatment and wing colour in the choice experiment. Shown are the results of a Chi-squared analysis of male mating success by male social treatment (no-rival or rival) or male wing colour (black or red).

**Figure 2.**
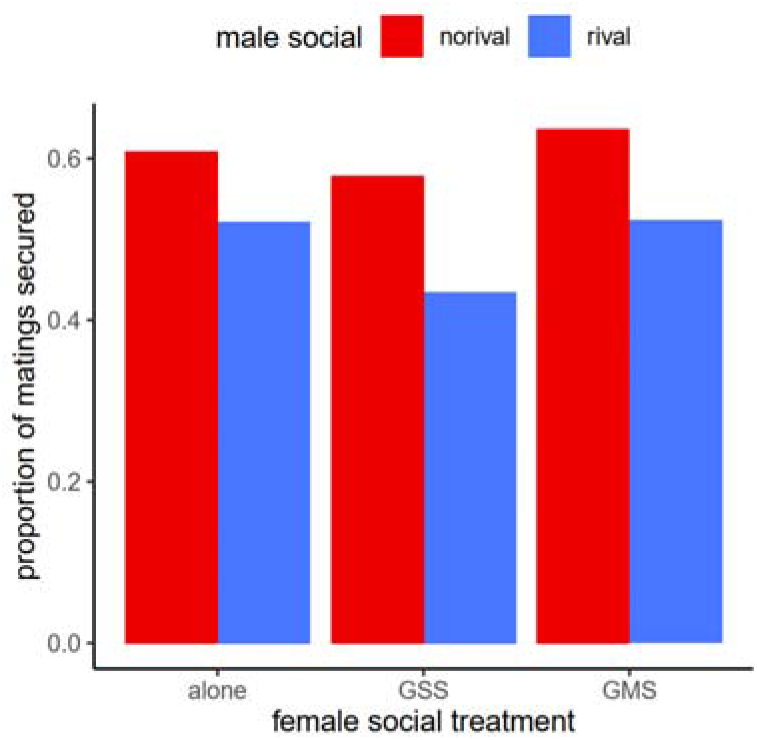
Proportion of matings secured by males from ‘no-rival’ and ‘rival’ male social treatments within each female treatment in the choice experiment. Results are shown for the 3 different female social treatments: alone, Group Same Sex (GSS), and Group Mixed Sex (GMS). Sample sizes in table S1.

**Figure 3.**
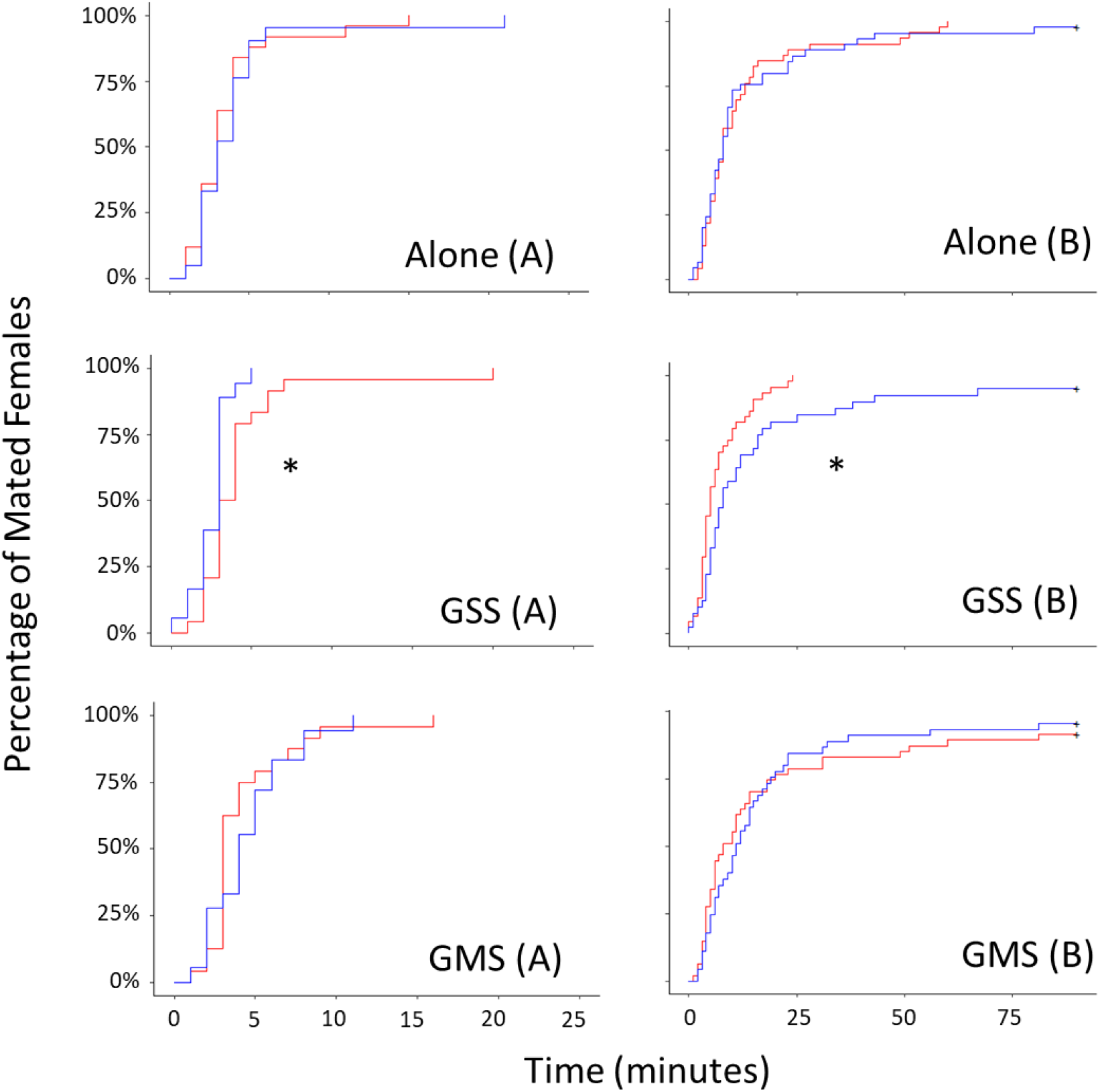
Mating latency of no-rival (red) and rival (blue) males with females from each of the female social treatments: Alone, Group Same Sex (GSS), Group Mixed Sex (GMS)) in the choice (A) and no-choice (B) experiments. Females and males used in these assays were raised in the different pre-mating social environments indicated (Figure 1A) for 48h prior to mating. Females were placed into a vial with both a rival and no-rival male in the choice experiment or either a rival or no-rival male in the no-choice experiment (figure 1B) and observed for 90 minutes for mating to begin and end. Matings occurred more rapidly in the choice experiment, hence note the different X axis in A vs B. Asterisks indicate significant differences between treatments (p<0.05). Sample sizes in table S1.

**Figure 4.**
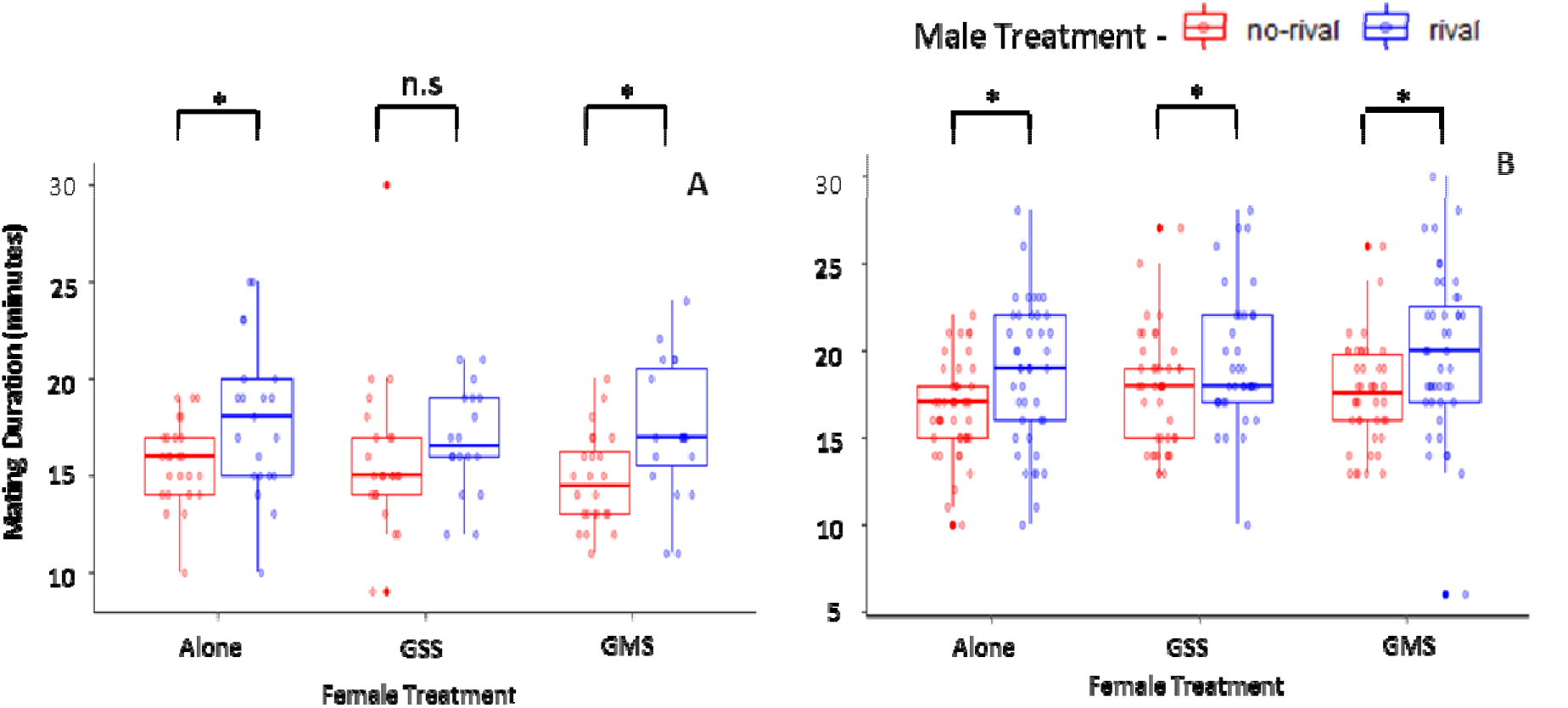
Mating duration (minutes) in the choice (A) and no-choice (B) experiments. Females and males used in these assays were raised in different pre-mating social environments for 48h prior to mating (Figure 1A). Female social environments: alone, group same sex (GSS), group mixed sex (GMS). Male social environments: no rival (red), rival (blue). Asterisks indicate significant differences between treatments (p<0.05). Shown are box plots (median, 25-75% IQ range, whiskers (1.5 x IQR) and outliers). Sample sizes in table S1.

**Figure 5.**
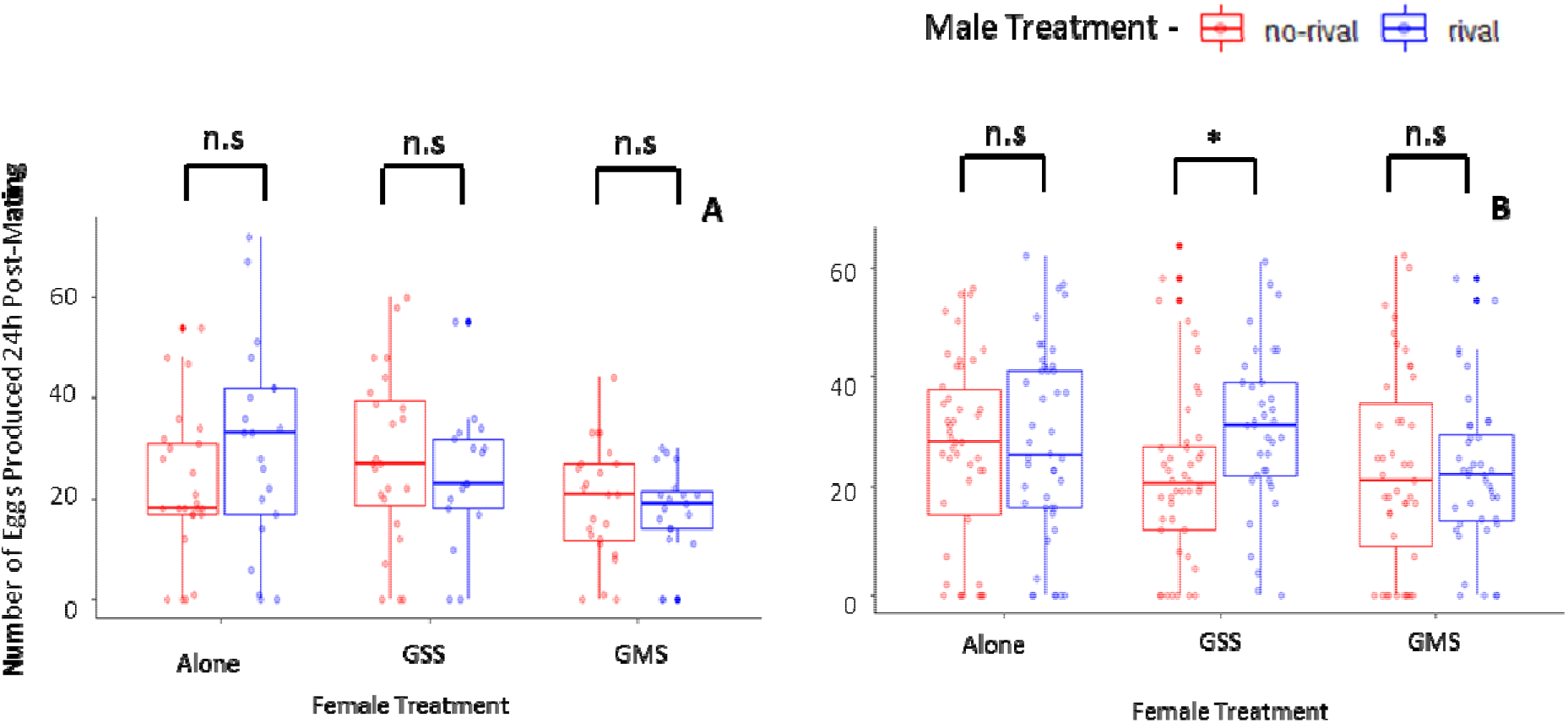
Number of eggs produced in the separate choice (A) and no-choice (B) experiments, in the 24h after mating. Females and males used in these assays were raised in different pre-mating social environments for 48h prior to mating (Figure 1A). Female social environments: ‘alone’, group same sex (GSS), group mixed sex (GMS). Male social environments: no rival (red), rival (blue). Asterisks indicate significant differences between treatments (p<0.05). Box plots as defined in figure 4. Sample sizes in table S1.

Overall, in these choice tests, the results showed evidence for interacting plasticity effects, but only for mating latency. Specifically, males previously exposed to rivals mated significantly faster only when paired with females previously exposed to other females (GSS). In contrast, mating duration was determined solely by the male’s social environment, being longer following exposure to rivals prior to mating. There was no effect of plasticity in either sex on fecundity.

### 2: Effect of male and female social environment on mating behaviour and fecundity under no-choice conditions

Here each female was presented with a single male in the mating arena. This tested the effect of the pre-mating social environment on mating behaviour and fecundity, in the absence of intrasexual competition between males in the mating arena. Mating latency was slower than in the above choice experiment, with a median mating latency of 8mins. Consistent with the choice trials, there were no significant differences in mating latency between rival and no-rival males paired with alone (*HR =* 0.965, *95% CI* [0.636, 1.466], *p =* 0.868) or GMS (*HR =* 1, *95%* CI [0.654, 1.53], *p =* 0.999) females (figure 3). As with the choice trials, mating latency was significantly different between male treatments for the GSS females, but the direction was reversed - rival males took significantly longer to start mating with GSS females than the no-rival males (*HR* = 0.519, *95% CI* [0.327, 0.825], *p =* 0.006). Females from all three treatments mated with rival males for significantly longer than no-rival males (alone: *t_86_ =* 2.831, *p =* 0.0058; GMS: *t_83_ =* 2.838, *p =* 0.0057; GSS: *t_78_* = 2.218, *p =* 0.0295) (figure 4B). As with the choice experiment, there was no difference in post-mating egg production following a mating with a rival and no-rival male in the alone and GMS female treatments (alone: *t_88_* = 0.368, *p =* 0.714; GMS: t_81_ = −0.181, *p =* 0.856). However, females in the GSS treatment that mated with rival males produced significantly more eggs than those that mated with no-rival males (t_78_ = 2.279, *p =* 0.0254) (figure 5B).

Overall, in these no-choice tests there was again evidence for effects of interacting plasticity – on mating latency, as above, but also upon fecundity. Interestingly, the mating latency effect was in the opposite direction, with males previously exposed to rivals being slower to mate with GSS, but not other treatment females. For fecundity, GSS, but not other treatment females laid significantly more eggs following matings with rival males. Again, mating duration was determined only by the male’s social environment, with rival males mating for longer across all contexts.

#### Overall patterns in choice and no-choice scenarios

The results from the two separate experiments above show evidence for interacting effects of plasticity, on mating latency across both scenarios and for fecundity in the no choice tests. The interactions pivoted around divergent responses seen with females from the GSS female social environment. However, we note potential limitations to the comparisons of the patterns obtained from the choice and the no choice tests. First, the experiments were done separately and second, there were differences in sample sizes (each male treatment in the choice experiment was approximately half that in the no-choice assays, as only half of the choice males secured a mating). To explore the latter, we assessed the potential for the differences in sample size to confound the patterns observed. Specifically, we considered whether the choice were underpowered in comparison to those of the choice assays. We did this in a resampling exercise (supplementary information) to explore the effect of sample size. The hypothesis tested was whether there was any evidence for not observing an effect in the choice assays that was present in the no-choice data, i.e. effects that ‘disappeared’ in the subsampled no-choice dataset.

For mating latency, the subsampled datasets showed broadly similar patterns and in no case was the direction of the effect reversed from the full no-choice dataset (figure S1, S2, S3). We conclude that the differences observed in both the choice and no-choice assays for mating latency were robust. For mating duration, in the full no-choice dataset we observed consistent significant differences between rival and no-rival males, and in the subsampled datasets showed significant differences in approximately half of the cases (figure S4). This was in line with the pattern of results seen in the choice data, where differences were either statistically significant or in the same direction. Hence, we conclude that the observation of male-only plasticity affecting mating duration across both test environments was reasonably robust. Finally, for fecundity, the observed difference between the male treatments in the no-choice assay interacting with the GSS females was not evident in any subsampled datasets (figure S5). This does not indicate that the no choice interacting plasticity effect was not robust, but could suggest that, had there been a comparable effect in the choice assays, it might not have been detected.

## Discussion

The major finding was that the interacting plastic responses of both sexes to their social and sexual environment can influence the expression of fitness-related traits. We found that mating latency and fecundity were sensitive to the interacting effects of responses to the socio-sexual environment made by both sexes. However, not all traits were affected in this way and we found that mating duration was determined largely by the social environment of the male. Effects of responses to the social environment by both sexes on mating latency were observed under both choice and no-choice conditions, but were manifested in opposing directions. Variation in the outcome of interacting plasticity pivoted around the outcomes with focal females previous exposed to other females (the GSS treatment). Our results show that the expression of some, but not all fitness-related reproductive traits can be determined by the outcome of interacting behavioural plasticity between the two sexes, and highlight the need for new predictive theory.

### Interacting phenotypic plasticity: mating behaviour and fecundity were determined by the plastic responses of both sexes

Importantly, the study provides evidence for the interacting effects of phenotypic plasticity expressed by both sexes on mating latency and fecundity. The direction of the interacting effect on mating latency as also reversed under choice versus no choice conditions. All of the interacting effects were centred around differences in the tests with the GSS, as opposed to other social treatment, females. This suggests that there was something qualitatively or quantitatively distinct about the plastic responses of these females, or the way they were perceived by males, in comparison to females kept on their own versus with males prior to mating. These findings are explored further, below.

In the no-choice experiment, no-rival males secured matings faster than did rival males with the GSS, but not other treatment, females. Mating latency of the rival vs no-rival males mating with GSS females in the choice experiment also differed significantly, although in this case no-rival males were slower to mate. The opposing direction of the response in the no choice vs choice experiments suggests the proximate mating environment as well as previous social experience both influence mating latency. It also suggests mating latency may be uncoupled from overall mate choice preference, since fewer rival males secured matings than no-rivals overall in the choice experiment, even though the rival males that did mate were faster to start copulating with GSS females. The design of mate choice experiments can affect preference outcomes, with females of many species generally exhibiting stronger preference when mates are presented simultaneously compared to sequentially (Dougherty and Shuker, 2014). Differences in preference between the designs could be driven by increased costs of rejection in a no-choice scenario (Dougherty and Shuker, 2014). It is possible that the outcomes we observed here could be a result of differential costs of rejection between the two mating regimes, combined with expectations of mate or resource competition from previous and proximate environments.

Overall, the differences in mating latency were focussed around the interactions with the female GSS treatment in both experiments. This suggests that the female socio-sexual environment can affect mating latency, but that variation in this trait is also influenced by interacting plasticity between both sexes. Given that the GSS females may perceive that mating opportunities are low, they may be less resistant to mating attempts than females from other treatments. The potentially lower choosiness of GSS females could exacerbate differences between rival and no-rival males in this treatment. Similarly, it was only females in the GSS treatment in the no-choice experiment that showed elevated fecundity when mating to rival, in comparison to no-rival, males. This implies that fecundity is influenced by both male and female pre-mating social environments. Female fecundity is affected by the receipt of SFPs. Processing of SP and ovulin is dependent on a network of female-expressed proteins and so there is opportunity for the female to exert control over the effects of SFPs (Sirot et al., 2015). Indeed, it may benefit the female to precisely control the level of SFP processing in response to her own perception of the chances of re-mating and the availability of resources such as nutrients and oviposition sites. For example, the costs of receiving SP are likely caused by a lower re-mating rate and increased short-term investment in egg production which may trade off against somatic investment. However, if opportunities for re-mating are low, as might be signalled in the GSS treatment when females are exposed to other females, then increased short-term fecundity mediated through SFPs may benefit females, or at least be costly to resist.

### Interacting plasticity and sexual conflict

The interactions described above between the behavioural plasticity expressed by each sex, have previously been investigated through theoretical modelling and are predicted to be an important facet of intra- and interspecific interaction dynamics (Yamaguchi and Iwaga, 2015; McLeod and Day, 2017; Day and McLeod, 2018). Specifically, sexual conflict is predicted under some circumstances to drive interacting plasticity to reach a fitness optimum between the interests of the male and the female, though the extent to which this occurs will depend upon the strength of sex-specific selection and thus whether either sex has the upper hand in any conflict.

We can consider whether there is any evidence to support this scenario in the findings we observed here. We saw that males that perceived themselves to be at high risk of sperm competition mated for significantly longer with females from all social environments. It has previously been observed that longer matings under these circumstances can transfer more cost-inducing SFPs (Wigby et al., 2011). Thus, the heightened SFP allocation by such males should be evident in plastic responses in females to resist SFP effects. This could be evident as reduced willingness to mate with such males (slow mating latency or lower mating success) potentially modified by the female’s own information on the likelihood of meeting any additional males (as signalled by their pre-mating and / or mating social environment). The results are generally in line with this expectation. Specifically, in the choice scenarios, rival males were generally less successful at mating than were no rival males and there were significant interactions of mating latency with female social status. Rival treatment males were significantly slower to mate with GSS females under no choice, but significantly faster under choice conditions. The data suggest that females may be able to assess their own social environment and respond in a manner that potentially mitigates SFP effects. We suggest that these findings indicated that selection may have favoured males that can increase their mating propensity while also favouring females that can effectively assess their environment. This would be interesting to investigate in future studies.

### Non-interacting phenotypic plasticity: mating duration was determined primarily by plasticity expressed by males

In contrast to the interacting effects described above, plasticity in mating duration was primarily determined by the responses of males to their social environments. In both the choice and no-choice assays, males of the rival treatment always mated for longer than males of the no-rival treatment (though not always significantly so). Therefore, we conclude the mating duration effect was independent of female social environment in both mating scenarios, indicative of a ‘one-sex’ plasticity exhibited by the male. This is consistent with previous findings (Lizé et al., 2011; Price et al., 2012; Bretman et al., 2013b). Under elevated sperm competition, mating duration may be correlated with an increased transfer of SFPs to the female (Bretman et al., 2009). As a result, females may experience reduced mating propensity (Mazzi et al., 2009). This is advantageous for the male, by preventing females from re-mating and thus increasing paternity share. However, these extended post-mating effects may also be costly to females if re-mating opportunities with higher quality males are lost. Plasticity in mating duration in response to rival exposure has previously been shown to be under male control (Bretman et al., 2013b). However, *Drosophila* females do have the ability to exert some influence on mating duration (Mazzi et al., 2009), and so the fact that the social environment of the female did not affect mating duration in our study suggests that the costs to males from sperm displacement are greater than costs to females from missed matings. Hence, we suggest there may be greater selection acting on males to guard females through increased mating duration than on females to resist longer matings.

### Non-interacting phenotypic plasticity: mate choice

In each of the three female social environment treatments within the choice experiment, no-rival males secured more matings than did rival males, although this was marginally non-significant. The potentially higher mating success of no-rival males over rival males could be due to female preference or to male competition. To respond to sperm competition, rival males increase the transfer of SFPs during mating (Wigby et al., 2009; Hopkins et al., 2019). No-rival males may be more attractive to females if they have remained in better condition. For example, rival treatment males may have experienced aggressive interactions resulting in physical damage (Davis et al., 2018). This could decrease their perception as high quality males or compromise their ability to court females (e.g. via wing damage). Such males could suffer reduced mating success (Chen, 2002; Davis et al., 2018). Alternatively, no-rival males may also secure more matings due to their ability to outcompete rival males. There are two possible explanations. First, territorial aggressive behaviour can occur between rival males in close proximity (Chen, 2002). This would be detrimental to their ability to compete for mates as they would have far less energy than the no-rival mate to successfully court the female. Second, rival males may be less willing to court due to the perception of high competition (Weir et al., 2011). Courtship behaviour is energetically costly (Bretman et al., 2013a; Cordts and Partridge, 1996) and rival males may benefit from withholding courtship until competition is reduced.

## Conclusions

We have provided the first experimental evidence for interacting behavioural plasticity in the model organism *Drosophila melanogaster. We* showed that males and females can plastically respond to their socio-sexual environment to influence the expression of mating duration, mating latency, and fecundity. These plastic responses, while induced to increase the fitness interests of each sex, interact in the case of mating latency and fecundity and may reflect the outcome of sexual conflict. Our findings suggest that studies of reproductive behaviour should carefully consider the socio-sexual environment of both males and females.

## Acknowledgements

For help with experiments we thank: Alice A Dore, Nathan McConnell and Aleksandra Łukasiewicz. This work was supported by the NERC (grant NE/R000891/1 to TC, AB and EKF).

## Author contributions

EF, AB and TC devised the experiments, EF and SL conducted the research, collected and analysed the data; EF, SL and TC wrote the paper; all authors contributed to the final draft.

## Data accessibility

The raw data are deposited in the DRYAD data depository, doi to be added in proof and raw data were made available for reviewers.

