## Supplemental Information for "Reproductive plasticity in both sexes interacts to determine mating behaviour and fecundity"

**Supplementary Information**

**Comparison of statistical power in the Choice and No-choice assays**

**Supplementary Tables**

**Table S1:** Sample sizes of the experimental set up and analyses of each treatment in the choice and no choice experiments.

**Supplementary Figures**

**Figure S1:** Three replicates of randomly sampled mating latency data in the alone female treatment from the no-choice experiment.

**Figure S2:** Three replicates of randomly sampled mating latency data in the group mixed sex (GMS) female treatment from the no-choice experiment.

**Figure S3:** Three replicates of randomly sampled mating latency data in the group same sex (GSS) female treatment from the no-choice experiment.

**Figure S4:** Three replicates of randomly sampled mating duration data from the no-choice experiment.

**Figure S5:** Three replicates of randomly sampled fecundity data from the no-choice experiment.

**Comparison of statistical power in the Choice and No-choice assays**

Three response variables were measured in the choice and no choice mating assays (mating latency, mating duration and female fecundity). For mating duration, rival males mated significantly longer than no-rival males in all female treatments for the no-choice assay, but for the choice assay this difference was not significant in the group same sex (GSS) female treatment. For mating latency, there were significant differences between no-rival and rival males for the GSS female treatment in both choice and no-choice assays, but in opposite directions. For fecundity, GSS females laid significantly more eggs after mating to rival males compared with no-rival males in the no-choice assay, but this was not significant in the choice assay. We tested here whether the inconsistencies across assays might be influenced by the sample size difference between them. In the choice assay, sample sizes for each male treatment were approximately half that in the no-choice assays, because only half of the males from each treatment were able to secure a mating. We tested the possibility that the choice assay was underpowered by randomly subsampling the no-choice data so n was equivalent to the choice dataset, and then re-running the analyses. We repeated this process to generate three replicates of subsampled data.

For mating latency, the subsampled datasets also showed inconsistency, but in no case was the direction of the effect reversed from the full dataset (figure S1, S2, S3). We conclude that the differences observed between choice and no-choice assays for mating latency are robust. For mating duration, the full dataset always returned significant differences between rival and no-rival males, the subsampled datasets showed significant differences in approximately half of the cases (figure S4). This was in line with the pattern of results seen in the choice data, where differences were either statistically significant or in the same direction. Hence, we conclude that the observation of male-only plasticity affecting mating duration across both test environments was reasonably robust. For fecundity, no significant differences were detected between male treatments in any of the subsampled datasets (figure S5). This does not indicate that the interacting plasticity effect was not robust, but could suggest that, had there been a comparable effect in the choice assays, it might not have been observed.

**Table S1**: Sample sizes of the initial number of flies set-up and the resulting samples sizes for the analyses of each variable (mating latency, mating duration and fecundity) in the choice and no-choice experiments.

|  | **rival (♂) alone (♀)** | **no-rival (♂) alone (♀)** | **rival (♂) group same sex, GSS (♀)** | **no-rival (♂) group same sex, GSS (♀)** | **rival (♂) group mixed sex, GMS (♀)** | **no-rival (♂) group mixed sex, GMS (♀)** |
| --- | --- | --- | --- | --- | --- | --- |
| **Choice Experiment** | | | | | | |
| **Set up** | 46 | | 42 | | 43 | |
| **Mating Latency** | 21 | 25 | 18 | 24 | 18 | 24 |
| **Mating Duration** | 21 | 25 | 18 | 23 | 19 | 24 |
| **Fecundity** | 21 | 25 | 17 | 24 | 19 | 24 |
| **No Choice Experiment** | | | | | | |
| **Set up** | 45 | 46 | 39 | 44 | 45 | 47 |
| **Mating Latency** | 45 | 45 | 39 | 44 | 45 | 47 |
| **Mating Duration** | 42 | 46 | 37 | 43 | 43 | 42 |
| **Fecundity** | 44 | 46 | 36 | 44 | 40 | 43 |

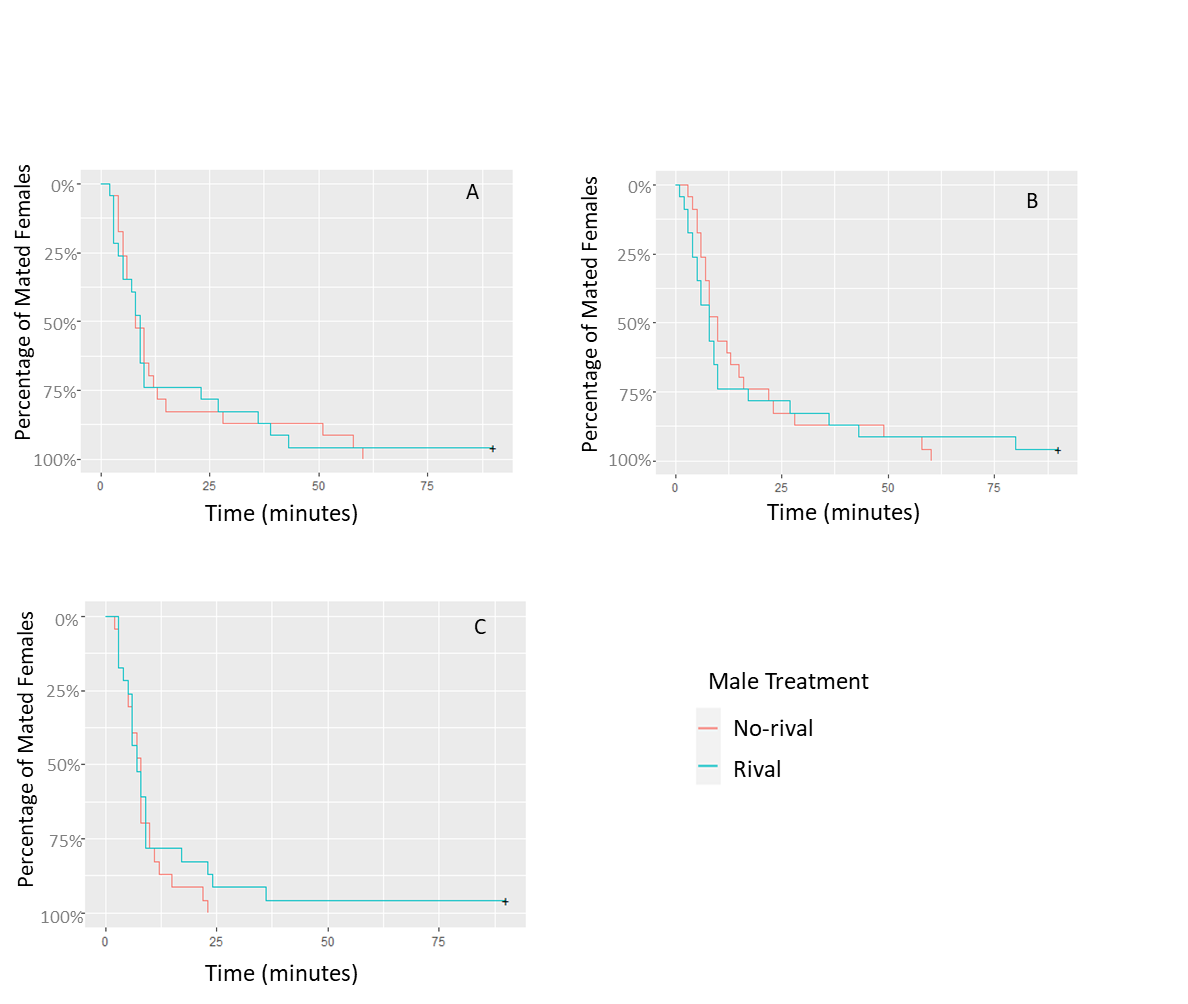

**Figure S1: Three replicates of randomly sampled mating latency data from the alone female treatment in the no-choice experiment.** The mating latency data (% of mated females against time in minutes) from the alone female treatment in the no-choice experiment were randomly sampled to give a sample size comparable to that of the choice experiment (rival: 23; no-rival: 23) and these data were then analysed using a GLM. Asterisks denote significant differences between rival and no-rival males (p<0.05). **A** – Hazard Ratio (HR)=1.06; 95% Confidence Intervals (CI) [0.589, 1.91]; p=0.845. **B** – HR=1.07; 95% CI [0.592, 1.947]; p=0.813. **C** – HR=0.769; 95% CI [0.418, 1.415]; p=0.399.

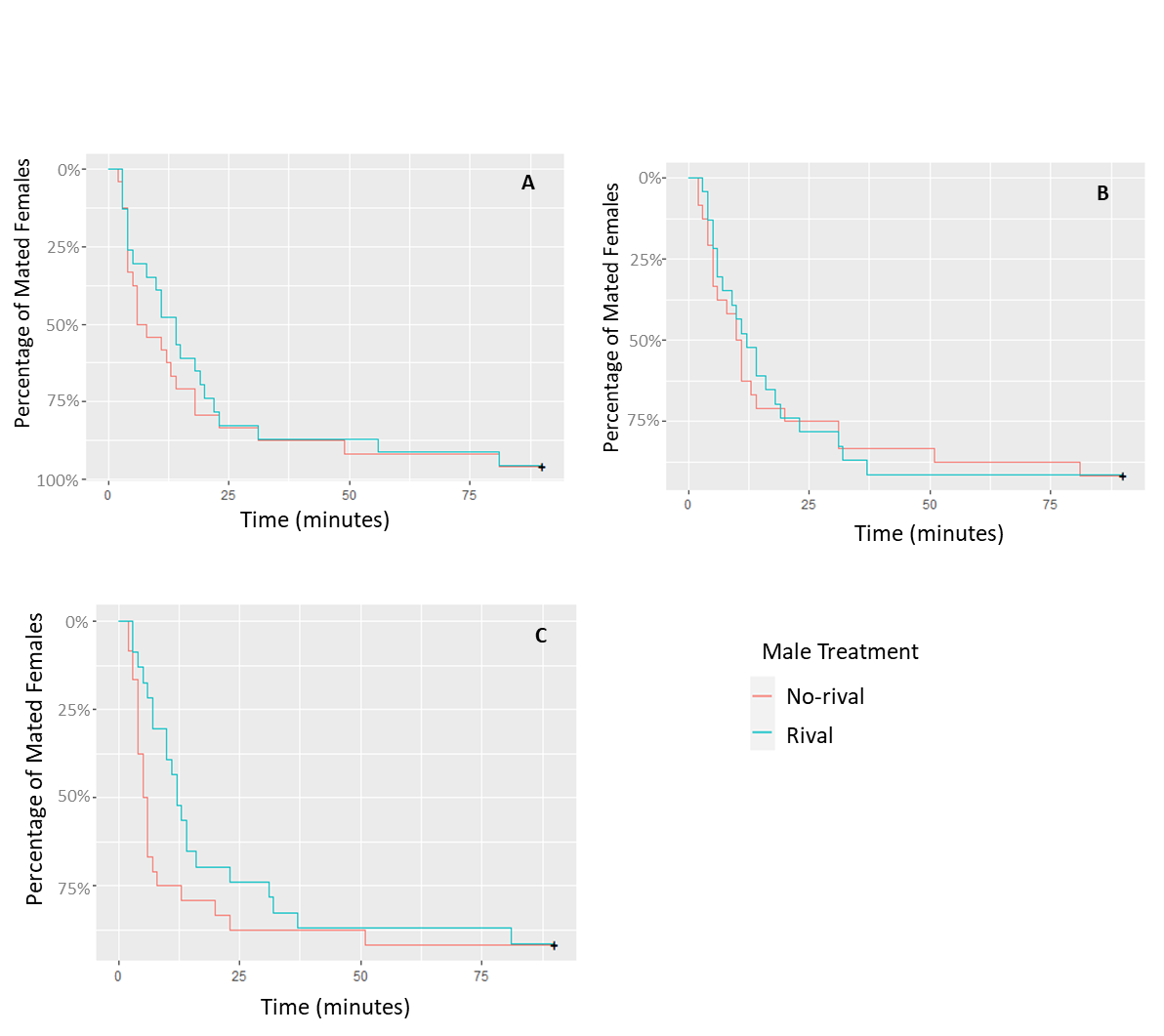

**Figure S2: Three replicates of randomly sampled mating latency data from the group missed sex (GMS) female treatment in the no-choice experiment**. The mating latency data (% of mated females against time in minutes) from the GMS female treatment in the no-choice experiment (rival: 23; no-rival: 24) were randomly subsampled to give a sample size comparable to that of the choice experiment and these data were then analysed using a GLM. Asterisks denote significant differences between rival and no-rival males (p<0.05). **A** – Hazard Ratio (HR)=0.832; 95% Confidence Intervals (CI) [0.463, 1.496]; p=0.539. **B** – HR=0.919; 95% CI [0.504, 1.675]; p=0.783. **C** – HR=0.610; 95% CI [0.333, 1.116]; p=0.109).

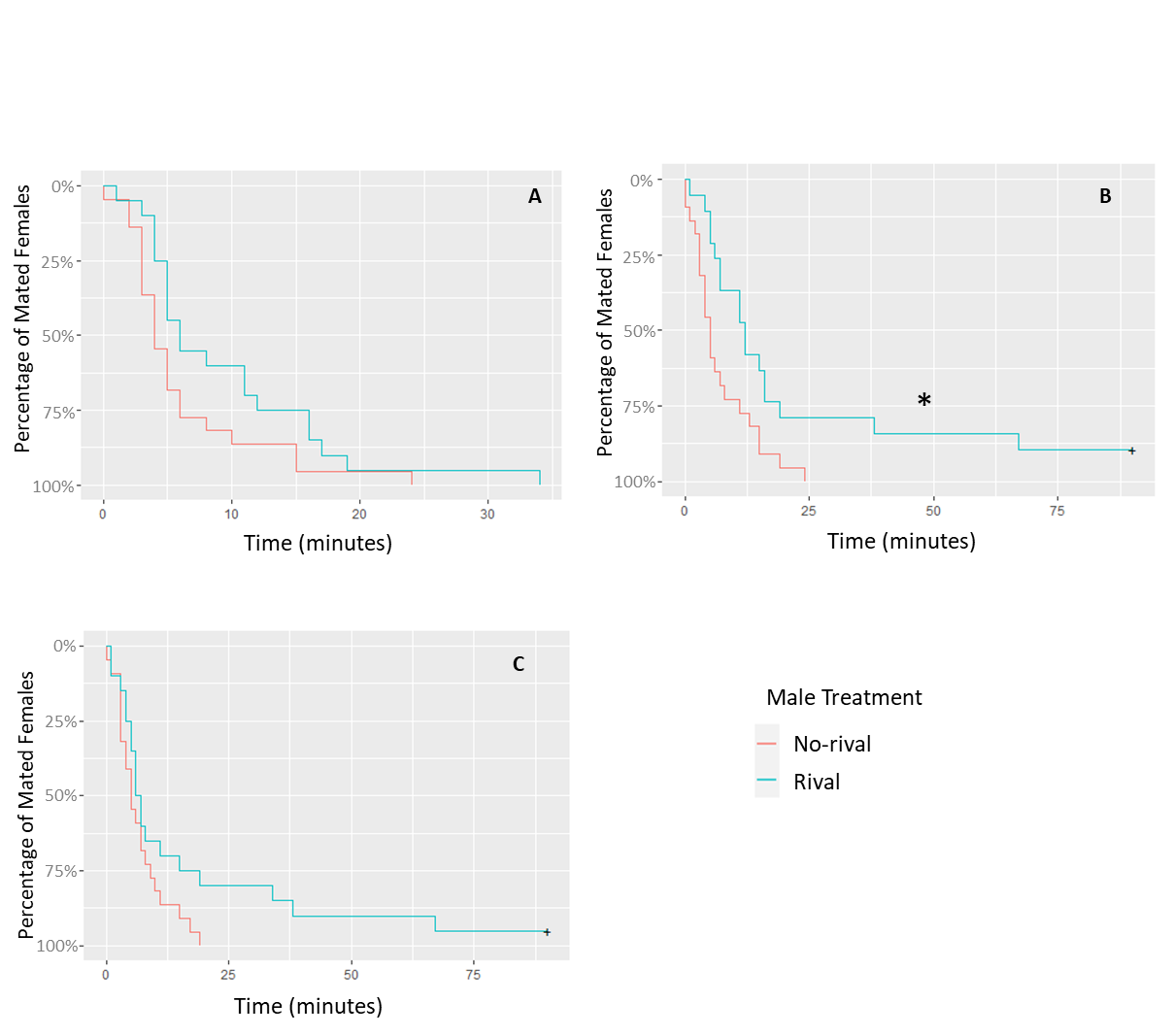

**Figure S3: Three replicates of randomly sampled mating latency data from the group same sex (GSS) female treatment in the no-choice experiment**. The mating latency data (% of mated females against time in minutes) from the GSS female treatment in the no-choice experiment were randomly subsampled to give a sample size comparable to that of the choice experiment (rival: 20; no-rival: 22) and these data were then analysed using a GLM. Asterisks denote significant differences between rival and no-rival males (p<0.05). **A** – Hazard Ratio (HR)=0.555; 95% Confidence Intervals (CI) [0.298, 1.037]; p=0.065. **B** – HR=0.413; 95% CI [0.212, 0.804]; p=0.009. **C** – HR=0.553; 95% CI [0.286, 1.069]; p=0.078.

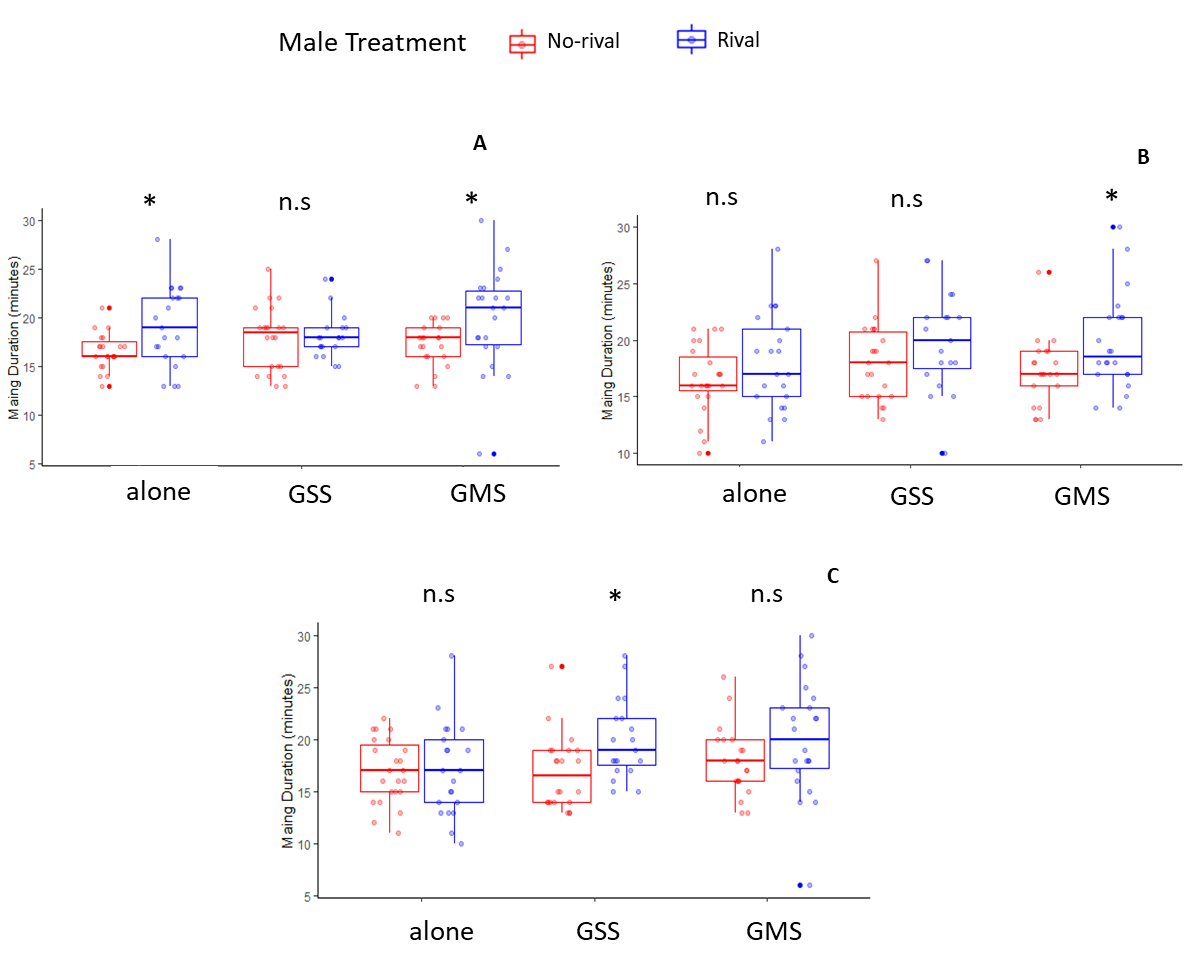

**Figure S4: Three replicates of randomly sampled mating duration data from the no-choice experiment.** The mating duration data (in mins) for each treatment from the no-choice experiment were randomly sampled to give a comparable sample size to that of the choice experiment (alone rival: 21; alone no-rival: 23; group rival: 19; group no-rival: 22; male rival: 22; male no-rival: 21) and these test data were then analysed using a GLM. Shown are box plots (median, 25-75% IQ range, whiskers (1.5 x IQR) and outliers). Asterisks denote significant differences between rival and no-rival males (p<0.05). **A** – alone: t=2.62, p=0.0122; group: t=0.262, p=0.795; male: t=2.295, p=0.0269. **B** – alone: t=1.303, p=0.2; group: t=1.502, p=0.141; male:t=2.285, p=0.0275. **C** – alone: t=0.122, p=0.903; group: t=2.708, p= 0.00999; male: t=1.538, p=0.132.

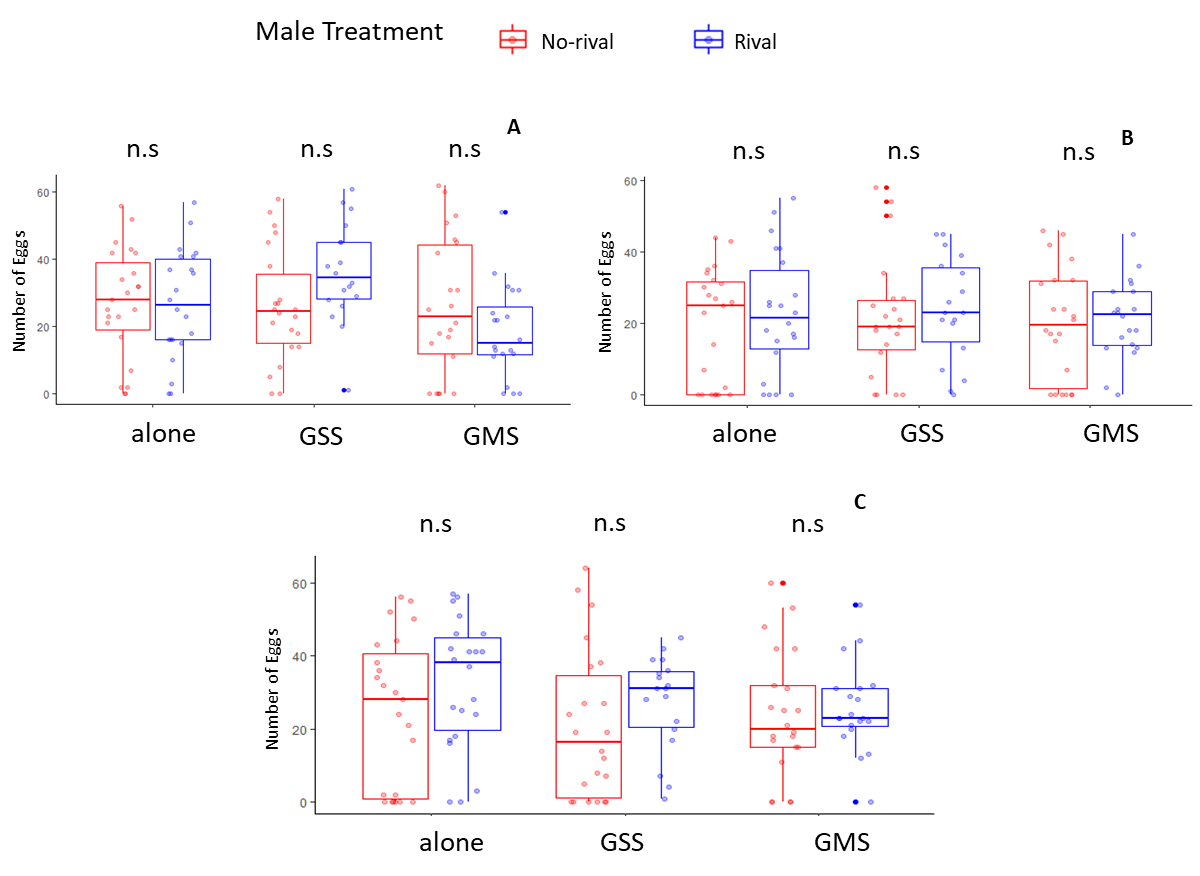

**Figure S5: Three replicates of randomly sampled fecundity data from the no-choice experiment.** The fecundity data for each treatment in the no-choice experiment were randomly sampled to give a sample size comparable to that of the choice experiment (alone rival: 22; alone no-rival: 23; group rival: 18; group no-rival: 22; male rival: 20; male no-rival: 22) and these data were then analysed using a GLM. Box plots as defined in figure S1. Asterisks denote significant differences between rival and no-rival males (p<0.05). **A** – alone: -t=0.039, p=0.969; group: t=1.939, p=0.0599; male: t=-1.419, p=0.164. **B** – alone: t=0.747, p=0.459; group: t=0.536, p=0.595; male: t=0.379, p=0.707. **C** – alone: t=1.316, p=0.195; group: t=1.12, p= 0.27; male: t=0.441, p=0.662.
